# Minimally-invasive nasal sampling in children offers accurate pneumococcal colonization detection

**DOI:** 10.1101/573196

**Authors:** Elissavet Nikolaou, Annie Blizard, Sherin Pojar, Elena Mitsi, Esther L. German, Jesús Reiné, Helen Hill, Paul S. McNamara, Andrea M. Collins, Daniela M. Ferreira, Simon P. Jochems

## Abstract

Nasopharyngeal colonization of potential respiratory pathogens such as *Streptococcus pneumoniae* is the major source of transmission and precursor of invasive disease. Swabbing deeply the nasopharynx, which is currently recommended by WHO, provides accurate pneumococcal detection but is unpleasant. We showed that nasal lining fluid filter strips offer equal detection sensitivity.

---

*Streptococcus pneumoniae* (Spn, pneumococcus), which naturally inhabits the nasopharynx of 40-95% of infants without causing disease [1] is one of the most frequent causes of bacterial infection in children. This bacterium accounts for about 38% of childhood deaths caused by pneumonia [2], which is the leading cause of death in children under 5 years worldwide [3]. Therefore, detection of pneumococcal colonization is of a great importance, as it is the primary reservoir for transmission and prerequisite of invasive disease.

There are a variety of sampling techniques for detecting nasopharyngeal colonization with different detection sensitivities. In adults, nasopharyngeal swab (NPS) and nasopharyngeal wash (NPW) cultures have been shown to detect higher rates of *S. pneumoniae* colonization than oropharyngeal swabs (OPS) [4]. However, sampling in children is challenging as swabs and aspirates can cause significant discomfort. Saliva sampling, which is painless to collect, has been successfully used to detect pneumococcus in children instead of NPS or OPS [5], however due to its polymicrobial nature might give false-positive results when using molecular methods [6].

On the other hand, sampling of nasal lining fluid using synthetic absorptive matrices (SAM) does not cause discomfort and has been used to detect respiratory syncytial virus (RSV) infection in a paediatric intensive care unit setting [7]. Whether such minimally-invasive samples could detect bacteria, including pneumococcus, has not been assessed yet, and there is a lack of evidence on whether nasal sampling is as sensitive as nasopharyngeal sampling for detection of carriage. The World Health Organisation (WHO) thus recommends NPS for pneumococcal colonization detection in children and both NPS and OPS in adults [8].

Recently, limitations of detection in conventional microbiology have led to the increased employment of PCR-based methods. The latter detects pneumococcus at low densities and thus offers high sensitivity for colonization detection. For detecting pneumococcal DNA in clinical samples, WHO recommends the use of quantitative PCR (qPCR) targeting the well-conserved autolysin-encoding gene *lytA* [8].

The present study aimed to test whether SAM can be used to accurately assess pneumococcal colonization by comparing the sensitivity (colonization rates and density) for detecting pneumococcal colonization in children between SAM and NPS using l*ytA* qPCR. We also compared the results obtained with NPS cultures.

## Methods

### Study design and ethics statement

SAM (Nasosorption™, Hunt Developments) and NPS (Transwab, Sigma) samples were collected from 49 children aged 1-5 years that were under general anaesthesia for unrelated reasons. Samples were collected after onset of general anaesthesia but prior to start of their planned procedure (dental extraction, MRI, orthopaedic or plastic surgery). To assess pneumococcal colonization, NPS samples were placed in skim milk, tryptone, glucose, and glycerine (STGG) medium and cultured on Columbia blood agar supplemented with 5% horse blood (PB0122A, Oxoid/Thermo Scientific) and 80μl gentamycin 1mg/mL (G1264-250mg, Sigma-Aldrich co Ltd). Plates were incubated overnight at 37°C and 5% CO_2_. Pneumococcal serotype was confirmed by latex agglutination (Statens Serum Institute, Copenhagen, Denmark). SAM samples and the remainder NPS samples were frozen at −80°C to be used for DNA extraction and qPCR.

Informed consent was obtained from all children’s parents after a thorough explanation of the study. This trial was approved by The National Health Service Research and Ethics Committee (REC) (17/NW/0663) and was sponsored by the Liverpool School of Tropical Medicine. All experiments were adapted to the relevant regulatory standards (Human Tissue Act, 2004).

### Pneumococcal DNA extraction from SAM and NPS samples

On the day of the extraction, SAM samples were thawed for 30 minutes on ice. 100μl of Luminex assay diluent (Thermofisher, Basingstoke, UK) was added to the filter and centrifuged at 1,503xg for 10 minutes at 4°C. After centrifugation, the eluted liquid was moved to a clean Eppendorf tube and centrifuged at 16,000xg for 10 minutes at 4°C. The supernatant was removed, and the pellets were used for DNA extraction. DNA extraction was performed using the Agowa Mag mini DNA extraction kit (LGC genomics, Berlin, Germany) and manufacturer’s instructions were folowed. For NPS samples, 200ul raw material were defrosted and DNA was extracted using the same procedure.

### Quantification of pneumococcal DNA in SAM and NPS samples by lytA qPCR

Colonization density in both SAM and NPS samples was determined by qPCR targeting the *lytA* gene (10) using the Mx3005P system (Agilent Technologies, Cheadle, UK). The sequences of the primers and probes used are: *lytA* forward primer: 5’-ACG-CAA-TCT-AGC-AGA-TGA-AGC-A-3’; *lytA* reverse primer 5’-TCG-TGC-GTT-TTA-ATT-CCA-GCT-3’; *lytA* probe: 5’-(FAM)-TGC-CGA-AAA-CGC-TTG-ATA-CAG-GGA-G-(BHQ-1)-3’. For the standard curve, pneumococcal DNA was extracted using the QIAamp DNA mini kit (Qiagen, Hilden, Germany). Samples were considered positive if two or all triplicates yielded a C_T_ < 40 cycles. Multiple experiment analysis was performed, and cross experiment threshold was calculated by using interrun calibrators.

### Statistical Analysis

Statistical analysis was performed by GraphPad Prism version 5.0 (California, USA). Data were log-transformed where appropriate. To distinguish between parametric and non-parametric data a Kolmogorov-Smirnoff normality test was performed. To quantify association between groups, the Pearson correlation test was used for parametric groups. Differences were considered significant if P ≤ 0.05.

## Results

### Both SAM and NPS qPCR detect equal pneumococcal colonization rates higher than NPS cultured samples

SAM and NPS samples were collected from 49/50 children enrolled in the study and used in this analysis. Using culture of NPS, 22/49 (44.90%) children were positive for Spn. Serotypes/groups identified were: SPN15 (7), SPN23 (4), SPN non-vaccine type (NVT) group G (3), SPN11 (2), SPN19 (2), SPN3, SPN10, SPN NVT group E and SPN NVT group I. All culture-positive children were also positive by molecular detection from NPS and SAM. Another 4 children were positive for Spn by *lytA* qPCR from both SAM and NPS. Moreover, another 5 children were positive by *lytA* qPCR from either SAM or NPS each. Thus, NPS and SAM *lytA* qPCR detected 31/49 (63.27%) children positive for Spn and agreed in 26/31 (83.87%) of them. In total, the number of positive colonised children detected by qPCR in both SAM and NPS samples was 1.4-fold higher than those detected in NPS cultured samples.

### Pneumococcal colonization densities measured by all three methods correlate significantly to each other

There was a significant correlation between bacterial load determined by SAM *lytA* qPCR, NPS *lytA* qPCR and NPS cultured (P<0.0001, Figure 1). In the majority of cases, pneumococcal densities measured by NPS qPCR were higher than those detected by NPS cultured (19/22, 86.36%). Four samples were positive by both SAM and NPS qPCR but not NPS cultured, with densities ranging between 10-176 DNA copies from SAM and 31-149 DNA copies from NPS. Another 5 samples were positive only by SAM *lytA* qPCR with densities 10-151 DNA copies. Another 5 were positive only by NPS *lytA* qPCR with densities 60-205 Spn DNA copies. Pneumococcal densities calculated by NPS qPCR were higher than those detected by SAM qPCR (24/26, 92.31%).

**Figure 1:**
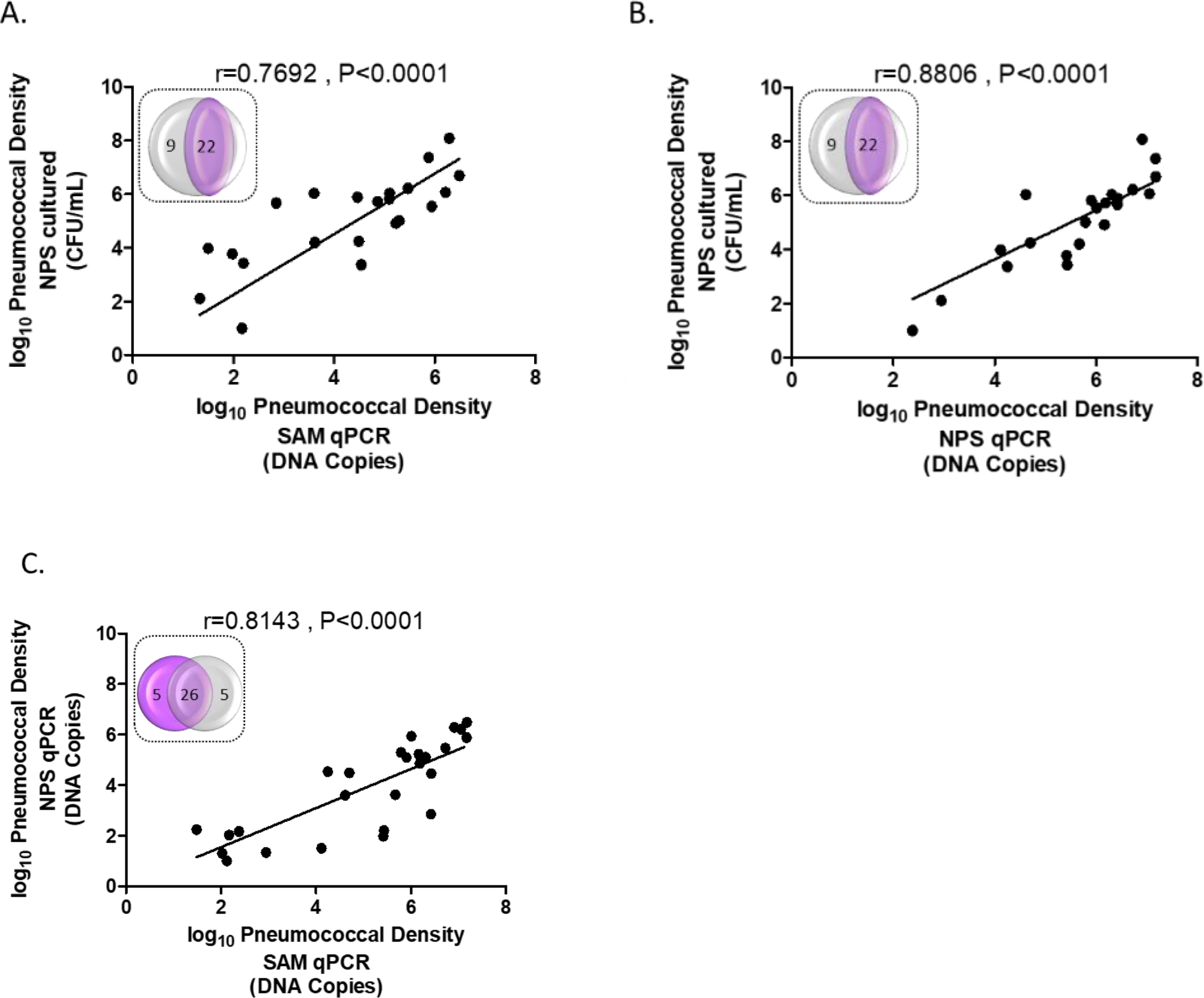
Correlation of colonization densities between detection methods A. SAM qPCR vs NPS cultured, B. NPS qPCR vs NPS cultured, C. SAM qPCR vs NPS qPCR. Points represent children positive for Spn and a linear regression line is added. Data was log transformed. A Pearson test was used to measure correlation between the methods of pneumococcal detection. Venn diagrams illustrate Spn positive children detected by each method; **A.** 22 children were positive for Spn by both SAM qPCR and NPS cultured (r=0.7692, ****P<0.0001). In total, SAM qPCR detected 31 Spn positive children (Venn diagram). **B.** 22 children were positive for Spn by both NPS qPCR and NPS cultured (r=0.8806, ****P<0.0001). In total, NPS qPCR detected 31 Spn positive children (Venn diagram), **C.** 26 children were positive for Spn by both SAM qPCR and NPS cultured (r=0.8143, ****P<0.0001). 5 children were Spn positive by *lytA* qPCR from either SAM or NPS each (Venn diagram).

## Discussion

Our results showed that SAM can be used as an alternative method to the current gold standard NPS [8] for pneumococcal detection in children with equal detection sensitivity. NPS sampling is associated with substantial discomfort [4]. SAM sampling targeting the anterior nares is a less invasive technique than NPS sampling, where a sample is collected from the nasopharynx. We have previously demonstrated that SAM sampling has low levels of discomfort, pain and lacrimation in adults [9].

The number of volunteers that were identified as Spn colonized by *lytA* qPCR (colonization rate) was higher than the number found by classical culture, as was expected. SAM qPCR detected equal numbers of Spn positive children as NPS qPCR (31/49, 63.27%) and agreed in 26/31 (83.87%) of cases, demonstrating that SAM sampling is a sensitive and specific alternative to NPS for pneumococcal detection in children. The children that were identified as carriers from only NPS or SAM were predominantly low density colonized and the discrepancy between the two sites might thus be stochastic. However, it is not impossible that differences between the two sites (anterior part of the nose and nasopharynx) exist in terms of microbiota composition. Additionally, we observed that pneumococcal densities in Spn positive volunteers detected by NPS qPCR are higher than those detected by SAM qPCR although this did not lead to differences in numbers of identified carriers. It is possible that swabbing collects more sample than absorption by SAM, however this did not affect sensitivity of Spn detection.

Previously, using the Experimental Human Pneumococcal Challenge (EHPC) model of infection in which healthy adults were challenged with 6B type pneumococcus, detection of pneumococcus in the nose of adults using SAM once Spn colonisation was established was low [10]. At day 2 and 6 after 6B exposure, only 1/9 (11.1%) and 1/7 (14%) Spn positive adults (carriers by classical culture of nasal washes) was found to be Spn positive by SAM qPCR. Possible explanations for this discrepancy are: differences in anatomy, physiology and nasal/oral microbiome between both groups and the possible change of colonization niche from the nasopharynx to oropharynx in adults [6]. The increased presence of pneumococcus in the anterior parts of the nose in children compared to adults could offer an explanation as to why children are transmitting more than adults.

In conclusion, our findings support that SAM sampling is a robust method for accurate detection of pneumococcus in children that could be employed during clinical trials and large epidemiological studies.

## Acknowledgements

We would like to thank all children for their participation and their parents for providing informed consent. Also, we would like to thank the Liverpool Alder Hey Children Hospital for supporting this research.

